# FIRST REPORT OF *CANDIDA AURIS* IN VIETNAM

**DOI:** 10.1101/2025.05.12.653631

**Authors:** Phu Truong Thien, Anh Nguyen Tuan, Tuan Nguyen Si

**Affiliations:** Cho Ray Hospital, Vietnam; Hong Bang International University

**Author notes:** Corresponding author: Anh Nguyen-Tuan | Cho Ray Hospital, Vietnam.

## Abstract

Early diagnosis and treatment of *Candida auris (C. auris)* infections improves mortality. At Cho Ray hospital, we successfully isolated and identified *C. auris* in all specimens by the Vitek 2 and Vitek MS systems. However, determining the sensitivity of *C. auris* is still difficult because currently neither the Clinical and Laboratory Standards Institute (CLSI) have standards for interpretation. At Cho Ray hospital, in order to provide clinical antifungal susceptibility testing for *C. auris*, the Department of Microbiology is relying on the provisional recommendations of the U.S. Centers for Disease Control and Prevention.

In Vietnam, there are currently no reports on the sensitivity of *C. auris*. In this article, we have the first report on the sensitivity of *C. auris* in Vietnam (specifically at Cho Ray Hospital). The method used in this study was the broth microdilution method recommended by the Clinical and Laboratory Standards Institute (CLSI), specifically using the Sensititre YeastOne antifungal panel. The Sensititre YeastOne antifungal susceptibility testing system (Thermo Fisher Scientific – USA) is based on the principle of broth microdilution combined with a colorimetric indicator, closely approximating the standard microdilution method. Each well contains a specific concentration of antifungal agent along with a color indicator. The minimum inhibitory concentration (MIC) is determined as the lowest concentration at which fungal growth is inhibited—indicated by the absence of a color change.

The Sensititre YeastOne panel offers several advantages: it is easy to perform, produces clear and interpretable results, and includes standardized guidelines for result interpretation, making it a convenient and reliable tool for antifungal susceptibility testing in clinical laboratories.

Result of the sensitivity rate (S) of *C. auris* to caspofungin was 81.6%, to fluconazole was 94.7%, and to amphotericin B was 71.1%. Resistance to all 3 antifungal groups (azole, echinocandins, polyene) was 0%, resistance to 2 antifungal groups accounted for 7.9% (3 out of 38 samples), resistance to 1 of 3 antifungal groups was 36.8%. At the same time, the similarity of 38 *C. auris* infections was also assessed using two methods: microdilution combined with color (Sensititre YeastOne – YO10) and diffusion according to concentration on agar plate (Etest) according to US CDC standards to help laboratories have more choices of methods suitable to the existing conditions in the laboratories.

## Introductionce

*Candida auris (C. auris)* is a new *Candida* agent, first announced in Japan in 2009 but to date more than 40 countries on all 5 continents have isolated this fungus [16]. *C. auris* is a yeast, growing with cells that can be single, in pairs or in groups. Cells are ovoid, elliptical or elongated with sizes ranging from 2.5 to 5 micrometers. *C. auris* rarely forms hyphae or pseudohyphae and also does not form germ tubes [27] but under special conditions, it can form fibers and change phenotypes between specific cell types [3] [23] [29] [33]. These traits may be related to virulence, fungal resistance, and the ability to survive adverse natural conditions and hosts. For example, when cultured on YTD medium (yeast extract, tryptone and dextrose) plus 10% NaCl at high temperature, pseudofilament-like forms can be produced [2] [11].

Similar to other *Candida* species, *C. auris* is an opportunistic pathogen and is primarily associated with critically ill and immunocompromised patients. Risk factors for *C. auris* include older age, diabetes, surgery and recently, the presence of indwelling medical devices (e.g., central venous catheters), immunosuppressed states, hemodialysis patients, neutropenia, chronic kidney disease, or extensive use of antibiotics and/or antifungal drugs [4] [18] [20] [24] [30]. Mortality rates from invasive infections associated with *C. auris* are relatively higher than those associated with other *Candida* species. Mortality rates associated with *C. auris* infection range from 30% to 72% [8] [12] [21] [32]. Early identification of *C. auris* and prompt treatment with appropriate antifungal regimens increases the chances of survival [28]

At Cho Ray hospital (first-line hospital - special class in the southern region, southern Vietnam), the first case of *C. auris* infection was detected in May 2020 and new infections continue to appear every year. As of December 31, 2023, the number of *C. auris* infections at Cho Ray Hospital is 38 cases.

To determine the sensitivity of *C. auris* at Cho Ray hospital, we conducted the study in the direction of descriptive cross-sectional research, location at Cho Ray hospital, the research object is *Candida auris* strain isolated and identified by Vitek 2, Vitek MS from patient specimens. The sample size for the study was collected from 2020 to the end of 2023 (n=38) based on CLSI M39-A4 sample collection standards [10]. The process is carried out by two methods: color - matched microdilution (Sensititre Yeastone – YO10) and concentration - based diffusion on agar plates (Etest). From the research data, we have collected and have the first report on the sensitivity of *C. auris* in Vietnam and at the same time evaluated the similarity of the two methods.

## Method

### Diffusion Method Using Etest Strips on Agar Plates

When the Etest strip is placed on the surface of Mueller-Hinton Agar (MHA) inoculated with the test fungal isolate, the antifungal agents at different concentrations on the strip rapidly diffuse into the MHA. After incubation for 18–24 hours, an elliptical inhibition zone symmetrical to the Etest strip appears, indicating the susceptibility of the fungus to the antifungal agent.

Prepare MHA medium, Etest strips, and sterile distilled water tubes. Use a sterile loop to collect fungal colonies and prepare a 0.5 McFarland suspension in the sterile distilled water tube. Use a sterile cotton swab to evenly spread the suspension over the surface of the MHA medium. Place the Etest strips onto the agar surface. Incubate the plates at 35°C for 48–72 hours. Read the results.

### Colorimetric Microdilution Method (Sensititre YeastOne - YO10)

The Sensititre YeastOne antifungal susceptibility test is based on the principle of colorimetric microdilution. Each well contains antifungal agents at appropriate dilution concentrations along with a colorimetric indicator. Results are determined by identifying the lowest concentration of the antifungal agent that inhibits fungal growth (indicated by the absence of color change).

Use a sterile loop to collect fungal colonies and prepare a 0.5 McFarland suspension in a tube containing sterile distilled water. Transfer 20 μl of the suspension into a tube containing 11 ml of YeastOne Broth to achieve a final concentration of 1.5– 8×10^3^ CFU/ml. Within 15 minutes of completing the previous step, dispense 100 μl of the suspension into each well of a 96-well microtiter plate. Seal the wells with the adhesive cover. Incubate the plate at 35°C in a non-CO_2_ incubator.

## Result

1. Susceptibility testing of *C. auris* by the concentration - diffusion method on agar plates (Etest)

With sample size n = 38, after performing the test using the concentration diffusion method on agar plates (Etest), the following results were obtained: with caspofungin, the number of strains still sensitive (S) was 33/38 strains, accounting for 84.8%; the number of non - sensitive strains (R) was 5 strains, accounting for 13.2%. With fluconazole, the number of strains still sensitive (S) was 36/38 strains, accounting for 94.7%; the number of non - sensitive strains (R) is 2 strains, accounting for 5.3%. With amphotericin B, the number of strains that were still sensitive (S) was 36/38 strains, accounting for 94.7%; the number of strains that were not sensitive (R) was 2 strains, accounting for 5.3%

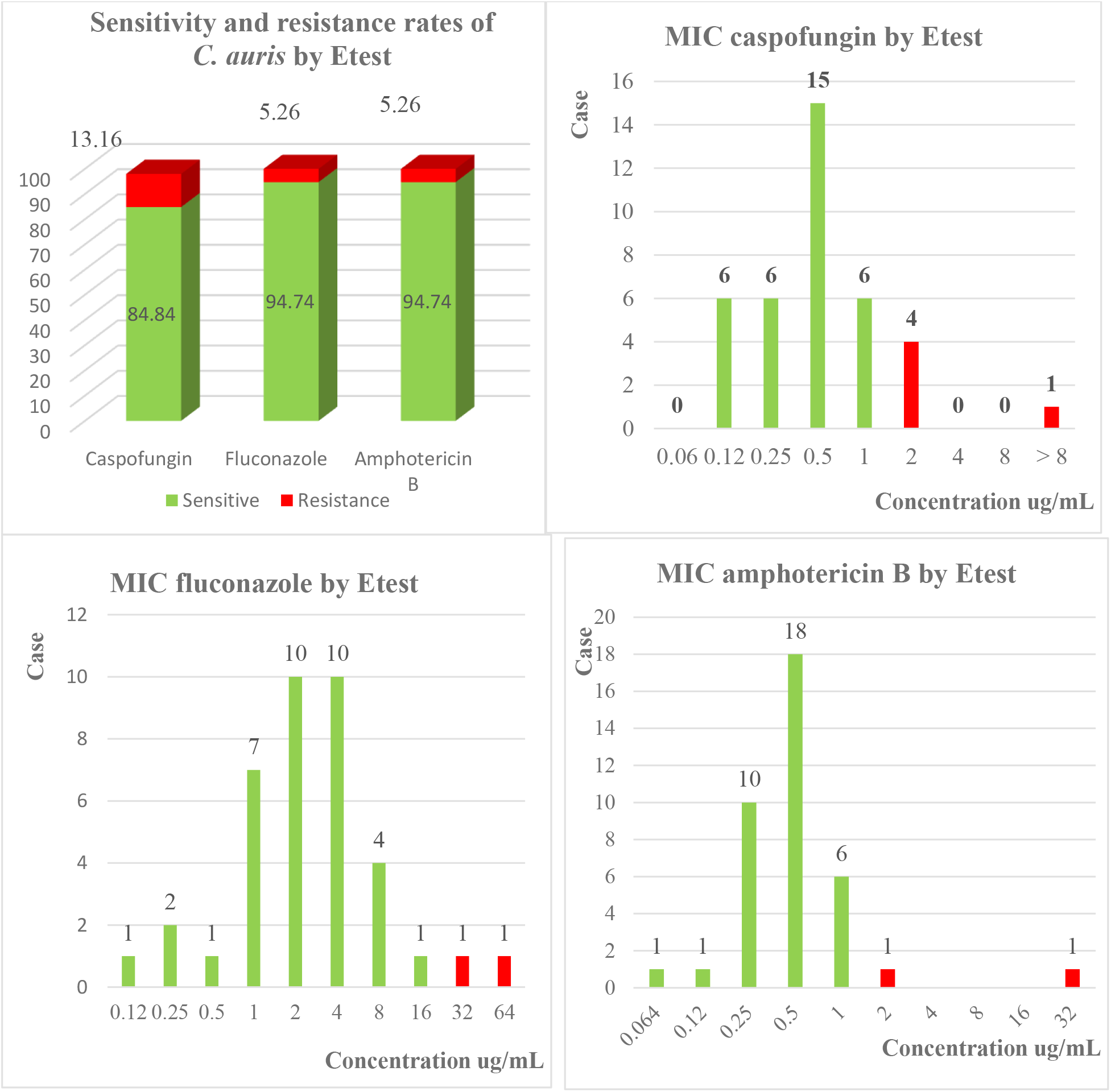

2. Susceptibility testing of *C. auris* by the colorimetric microphase method (Sensititre Yeastone - YO10)

In this study, the microdilution method was performed and the results obtained were: with caspofungin, the number of strains that were still sensitive (S) was 31/38 strains, accounting for 81.6%, the number of strains that were not sensitive (R) was 7 strains, accounting for 18.4%. With fluconazole, the number of strains that were still sensitive (S) was 36/38, accounting for 94.7%, the number of strains that were not sensitive (R) was 2, accounting for 5.3%. With amphotericin B, the number of strains that were still sensitive (S) was 27/38, accounting for 71.1%, the number of strains that were not sensitive (R) was 11, accounting for 28.9%.

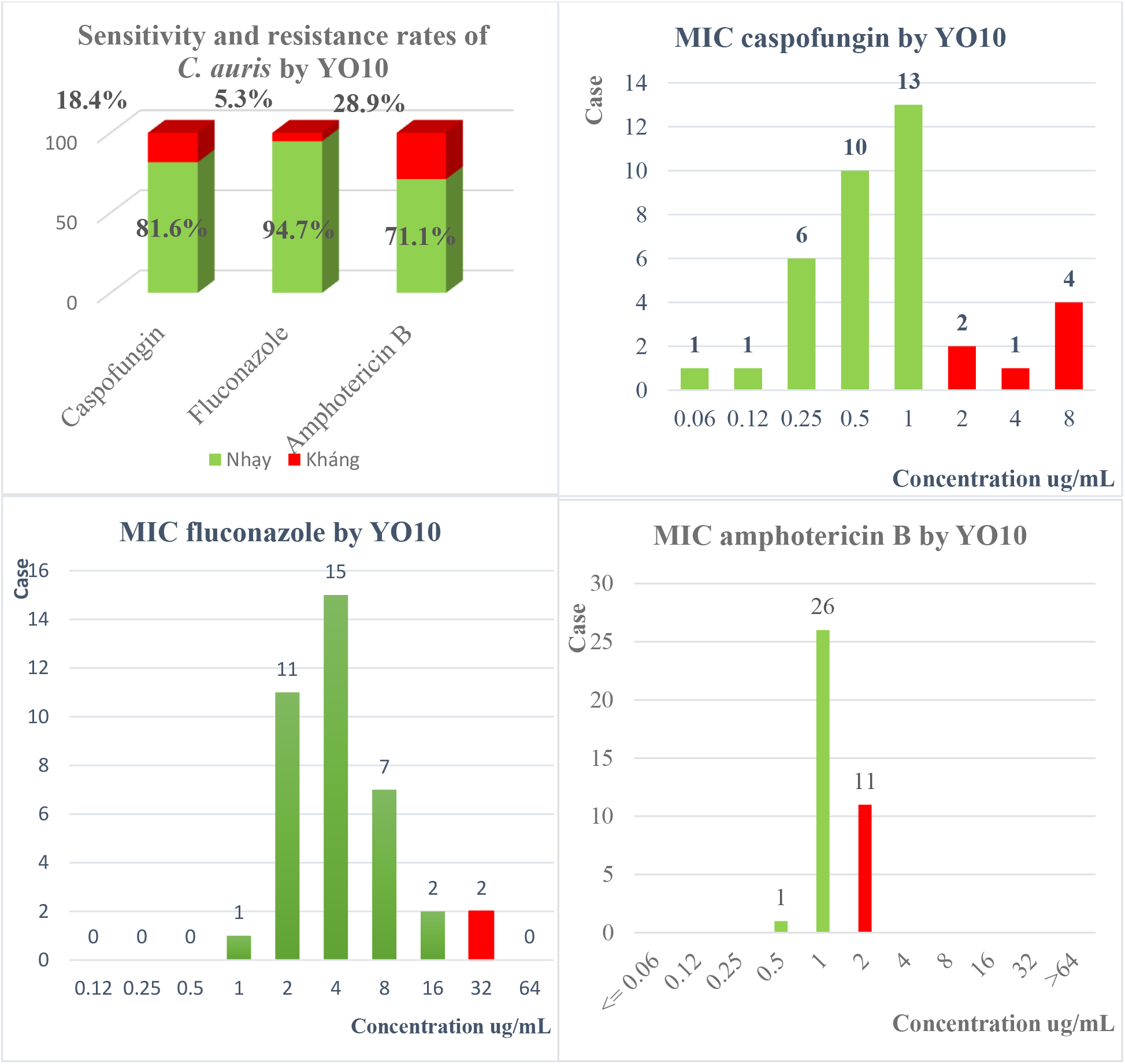

3. Similarity of susceptibility, resistance and MIC phenotypes of antifungals by Etest and YO10 methods

3.1. Similarity of caspofungin sensitivity, resistance and MIC phenotypes by Etest and YO10 methods

**Table 3.3:** Phenotypic results of caspofungin sensitivity and resistance between YO10 and Etest (n=38)

| Etest \ YO10 | Sensitive (S) |  | Resistance (R) |  |
| --- | --- | --- | --- | --- |
|  | Number | Ratio | Number | Ratio |
| Sensitive (S) | 31 | 81.6% | 2 | 5.3% |
| Resistance (R) | 0 | 0% | 5 | 13.1% |

Phenotypic similarity in sensitivity and resistance was 94.7%.
Phenotypic dissimilarity in sensitivity and resistance was 5.3%.

**Table 3.4:** Comparison of MIC results between YO10 and Etest on caspofungin (n=38)

| YO10 \ Etest | 0.06 | 0.12 | 0.25 | 0.5 | 1 | 2 | 4 | 8 | >8 |
| --- | --- | --- | --- | --- | --- | --- | --- | --- | --- |
| 0.06 |  |  |  |  |  |  |  |  |  |
| 0.12 | 1 |  | 3 | 1 | 1 |  |  |  |  |
| 0.25 |  |  | 2 | 2 | 2 |  |  |  |  |
| 0.5 |  |  | 1 | 5 | 7 | 2 |  |  |  |
| 1 |  | 1 |  | 2 | 3 |  |  |  |  |
| 2 |  |  |  |  |  |  | 1 | 3 |  |
| 4 |  |  |  |  |  |  |  |  |  |
| 8 |  |  |  |  |  |  |  |  |  |
| >8 |  |  |  |  |  |  |  | 1 |  |

Similar MICs
MICs are similar within $\pm 2$ dilutions
Mismatch MICs

3.2. Similarity of fluconazole sensitivity, resistance and MIC phenotypes by Etest and YO10 methods

**Table 3.5:** Phenotypic results of fluconazole sensitivity and resistance between YO10 and Etest (n=38)

| Etest \ YO10 | Sensitive (S) |  | Resistance (R) |  |
| --- | --- | --- | --- | --- |
|  | Number | Ratio | Number | Tỉ lệ |
| Sensitive (S) | 36 | 94.7% | 0 | 0% |
| Resistance (R) | 0 | 0% | 2 | 5.3% |

Phenotypic similarity in sensitivity and resistance was 100%
Phenotypic dissimilarity in sensitivity and resistance was 0%

**Table 3.6:** Comparison of MIC results between YO10 and Etest on fluconazole (n=38)

| YO10 \ Etest | 0.12 | 0.25 | 0.5 | 1 | 2 | 4 | 8 | 16 | 32 | 64 |
| --- | --- | --- | --- | --- | --- | --- | --- | --- | --- | --- |
| 0.12 |  |  |  |  |  |  | 1 |  |  |  |
| 0.25 |  |  |  |  | 1 | 1 |  |  |  |  |
| 0.5 |  |  |  | 1 |  |  |  |  |  |  |
| 1 |  |  |  |  | 3 | 2 | 2 |  |  |  |
| 2 |  |  |  |  | 4 | 4 | 2 |  |  |  |
| 4 |  |  |  |  | 2 | 7 | 1 |  |  |  |
| 8 |  |  |  |  | 1 | 1 | 1 | 1 |  |  |
| 16 |  |  |  |  |  |  |  | 1 |  |  |
| 32 |  |  |  |  |  |  |  |  | 1 |  |
| 64 |  |  |  |  |  |  |  |  | 1 |  |

Similar MICs
MICs are similar within $\pm 2$ dilutions
Mismatch MICs

3.3. Similarity of amphotericin B sensitivity, resistance and MIC phenotypes by Etest and YO10 methods

**Table 3.5:** Phenotypic results of amphotericin B sensitivity and resistance between YO10 and Etest (n=38)

| YO10 \ Etest | Sensitive (S) |  | Resistance (R) |  |
| --- | --- | --- | --- | --- |
|  | Number | Ratio | Number | Tỉ lệ |
| Sensitive (S) | 27 | 71.1% | 9 | 23.6% |
| Resistance (R) | 0 | 0% | 2 | 5.3% |

Phenotypic similarity in sensitivity and resistance was 76.4%
Phenotypic dissimilarity in sensitivity and resistance was 23.6%

**Table 3.8:** Comparison of MIC results between YO10 and Etest on amphotericin B(n=38)

| YO10 \ Etest | 0.064 | 0.12 | 0.25 | 0.5 | 1 | 2 | 4 | 8 | 16 | 32 |
| --- | --- | --- | --- | --- | --- | --- | --- | --- | --- | --- |
| 0.064 |  |  |  |  | 1 |  |  |  |  |  |
| 0.12 |  |  |  | 1 |  |  |  |  |  |  |
| 0.25 |  |  |  |  | 10 |  |  |  |  |  |
| 0.5 |  |  |  |  | 14 | 4 |  |  |  |  |
| 1 |  |  |  |  | 1 | 5 |  |  |  |  |
| 2 |  |  |  |  |  | 1 |  |  |  |  |
| 4 |  |  |  |  |  |  |  |  |  |  |
| 8 |  |  |  |  |  |  |  |  |  |  |
| 16 |  |  |  |  |  |  |  |  |  |  |
| 32 |  |  |  |  |  | 1 |  |  |  |  |

Similar MICs
MICs are similar within $\pm 2$ dilutions
Mismatch MICs

## Discuss

### 4.1. Evaluation of the similarity between Etest and YO10 on caspofungin antifungal

Echinocandins (specifically caspofungin) are considered first-line therapy for invasive fungal infections, although many studies have reported resistant isolates across different geographical regions, with the highest levels of resistance reported in India [5] [25]. Echinocandin resistance is often associated with hotspot mutations in the FKS1 gene, which encodes the enzyme 1-3-β-D-glucan synthase that is targeted by echinocandins, resulting in lower affinity of the enzyme for the drug [5]. A study by Kordalewska et al. identified that the S639F mutation in FKS1 conferred echinocandin resistance in four isolates from India [13], with another study identifying another amino acid substitution at the same position (S639P) [1].

**Figure 4.1:**
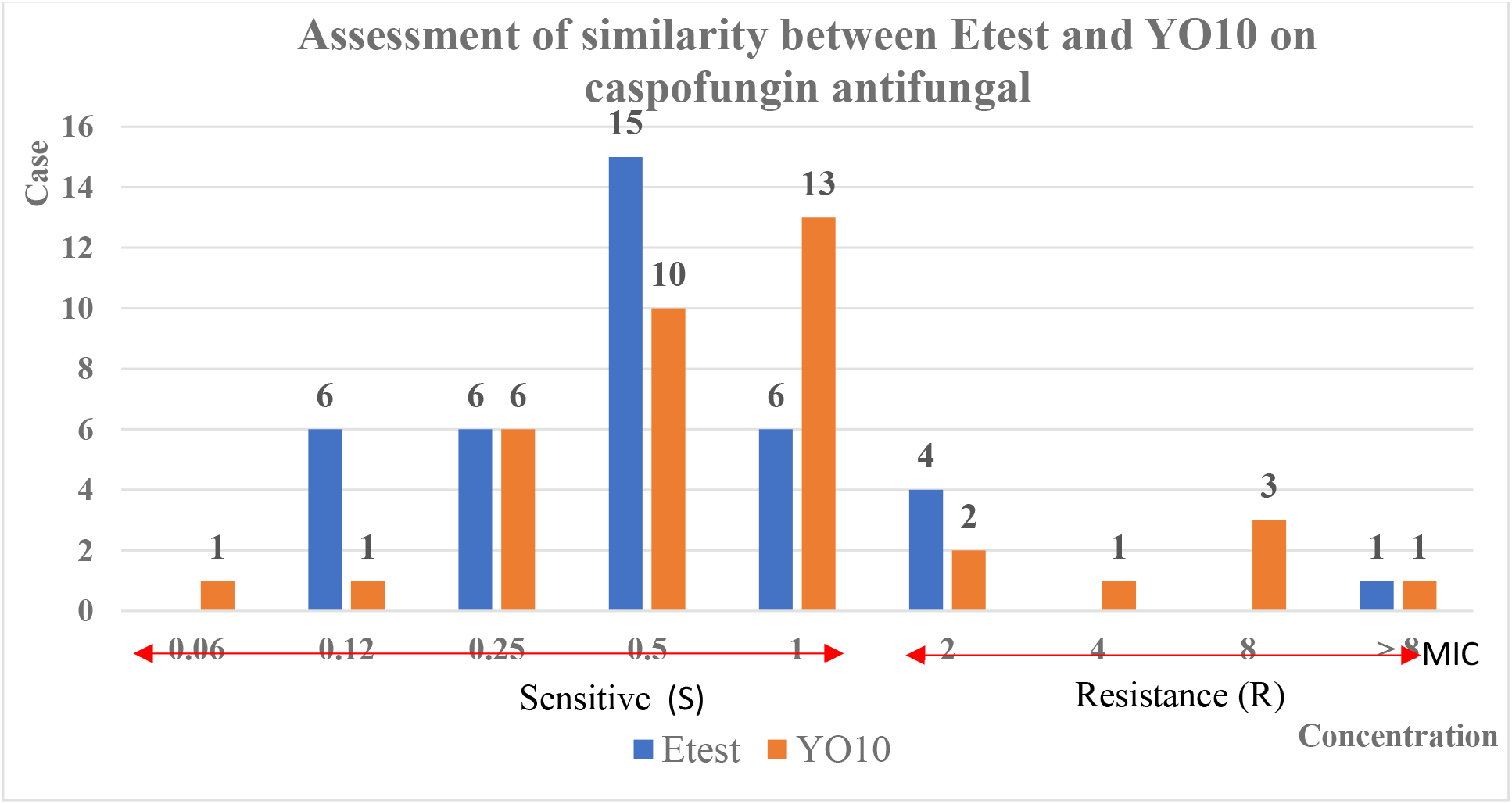
Assessment of similarity between Etest and YO10 on caspofungin antifungal

According to US CDC standards [22], breakpoint of *C. auris* on caspofungin ≥ 2 μg/ml was considered resistant. In this study, we found that both methods had similar MICs with 10 samples accounting for 26.3%. MIC within ± 2 dilution limits between the 2 methods was 26/38 samples, accounting for 68.4%. MIC discordance between the 2 methods was 2/38 samples, accounting for 5.3%. The MICs ranged from 0.25 – 1 μg/ml (MIC = 0.25 μg/ml in 6 samples, MIC = 0.5 μg/ml in 10 samples, MIC = 1 μg/ml in 6 samples; total 22 samples). Similarly, in terms of phenotypic sensitivity and resistance, the results of the antifungal test were similar in phenotype between the two methods in 36/38 samples, accounting for 94.7%, The results of antifungal tests were not consistent in phenotype between the two methods in 2/38 samples, accounting for 5.3%. MIC distribution ranged from 0.25 - 1 μg/ml (with Etest being 27 samples, YO10 being 29 samples). In addition, based on the YO10 antifungal tray, we obtained a caspofungin resistance rate of 18.4%, which is higher than the data from the US CDC (<2%) [22] as well as higher than the similar report in India (3.8%) [31]. The rate of caspofungin resistance in the Indian study was explained by the patients having a history of previous exposure to echinocandins [31] This may explain the high resistance in Vietnam, as hospitals use and recommend echinocandins (caspofungin) as first-line treatment. Additionally, during the testing, 4 test samples (1 MIC = 4 μg/ml, 3 MIC = 8 μg/ml) exhibited the Eagle effect (also known as the paradoxical growth effect). This phenomenon refers to the decrease in fungicidal activity at high concentrations of antifungal agents (when the concentration of antifungal agent is increased beyond a certain value, an increase in fungal survival is observed). This can be explained by the fact that at very high concentrations the molecules can compete with each other for binding to their receptors, for example in the case of penicillin the penicillin-binding protein) mainly in vitro [14]. Here, the paradoxical effect explanation becomes relevant, since high in vitro concentrations have not been shown to correlate with in vivo concentrations with isolates not carrying the FKS1 mutation [13]. From the above two reasons, we can see that the rate of resistance to caspofungin in Vietnam will be higher than in other regions and we can also see the similarity between the two methods in terms of sensitivity - resistance phenotype is 95% (36/38 samples).

### 4.2 Evaluation of the similarity between Etest and YO10 on fluconazole antifungal

Azole resistance mechanisms in *C. auris* are diverse and closely resemble those reported in other *Candida spp*, including mutations in the azole target gene (ERG11) and activation of the drug transporters (efflux pumps) CDR1 and MDR1 under the control of their respective transcription factors TAC1b and MRR1. Several gain-of-function mutations of TAC1b and MRR1 that confer azole resistance in *C. auris* have been reported previously [5].

According to US CDC standards [22], the breakpoint of *C. auris* on fluconazole ≥ 32 μg/ml is resistance.

**Figure 4.2:**
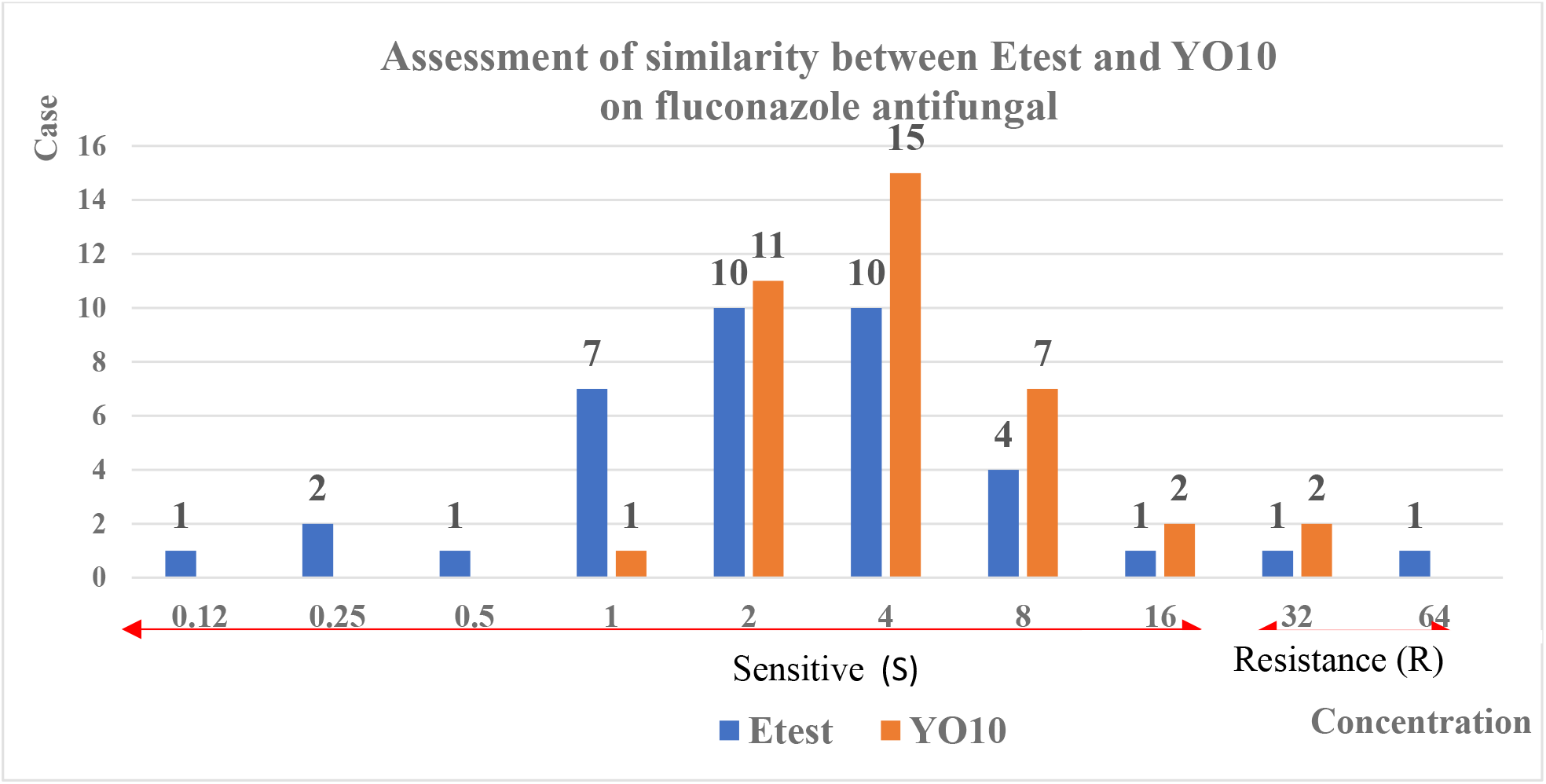
Assessment of similarity between Etest and YO10 on fluconazole antifungal

The results collected from the study showed that the similarity in MIC between the two methods was 36.7% (14 samples), MIC within ± 2 dilution limits between the 2 methods was 19/38 samples, accounting for 50%. MIC discordance between the 2 methods was 5/38 samples, accounting for 13.3%. MIC ranged from 2 – 8 μg/ml (MIC = 2 μg/ml had 10 samples, MIC = 4 μg/ml had 10 samples, MIC = 8 μg/ml had 4 samples, total of 24 samples). In terms of sensitivity and resistance phenotype, MICs were mainly distributed between 2 and 8 μg/ml and had the highest similarity among the 3 antifungals studied (100%). The results of the antifungal test were similar in phenotype between the two methods in 38/38 samples (with 36 sensitive samples and 2 resistant samples), accounting for 100%. The results of the antifungal test were not similar in MIC between the two methods in 0/38 samples, accounting for 0%.

According to the US CDC, the fluconazole resistance rate in the US is about 90% [22] but in Vietnam, the fluconazole resistance rate is only 5.3% (2 samples). This may be explained by the fact that these *C. auris* strains were almost the first strains discovered in Vietnam, so they have not yet shown much resistance to antifungal drugs, including fluconazole. In addition, studies are needed to determine whether the genome of *C. auris* circulating in Vietnam is truly similar to the genome of *C. auris* strains circulating in other areas.

### 4.3 Evaluation of the similarity between Etest and YO10 on amphotericin B antifungal

Amphotericin B (AmB) is a polyene antifungal drug approved in 1958. It is the first choice for the treatment of meningitis caused by *cryptococcus, mucorales*,*…* as well as infections caused by many dimorphic fungi. [15] Amphotericin B is a fungicide that, despite its toxic side effects on the liver and kidneys, remains the drug of choice for treating drug-resistant fungal infections, including those caused by *C. auris* [26]. The most widely accepted mechanism of action of AmB suggests that AmB may bind to or trap membrane ergosterol, trigger membrane permeability changes by sequestering ergosterol, or modulate channel function [7]. Although the exact mechanism of AmB action remains unclear and controversial, the general membrane dysfunction seems plausible, suggesting that AmB triggers “pore” formation in lipid membranes after binding or sequestering ergosterol [6], changes in membrane permeability lead to the formation of small “pores” and the disruption of the electrolyte gradient, causing the escape of intracellular potassium ions (K +) and thereby triggering cell lysis and death [19]. Because there are many hypotheses about the mechanisms of action of AmB such as how AmB can pass through rigid carbohydrate cell wall structures and how it inserts into lipid membranes to form “pore” structures, we are still far from fully understanding the mechanisms of action of AmB on *C. auris*.

According to US CDC standards [22], The breakpoint of *C. auris* on amphotericin B ≥ 2 μg/ml is resistance.

**Figure 4.3:**
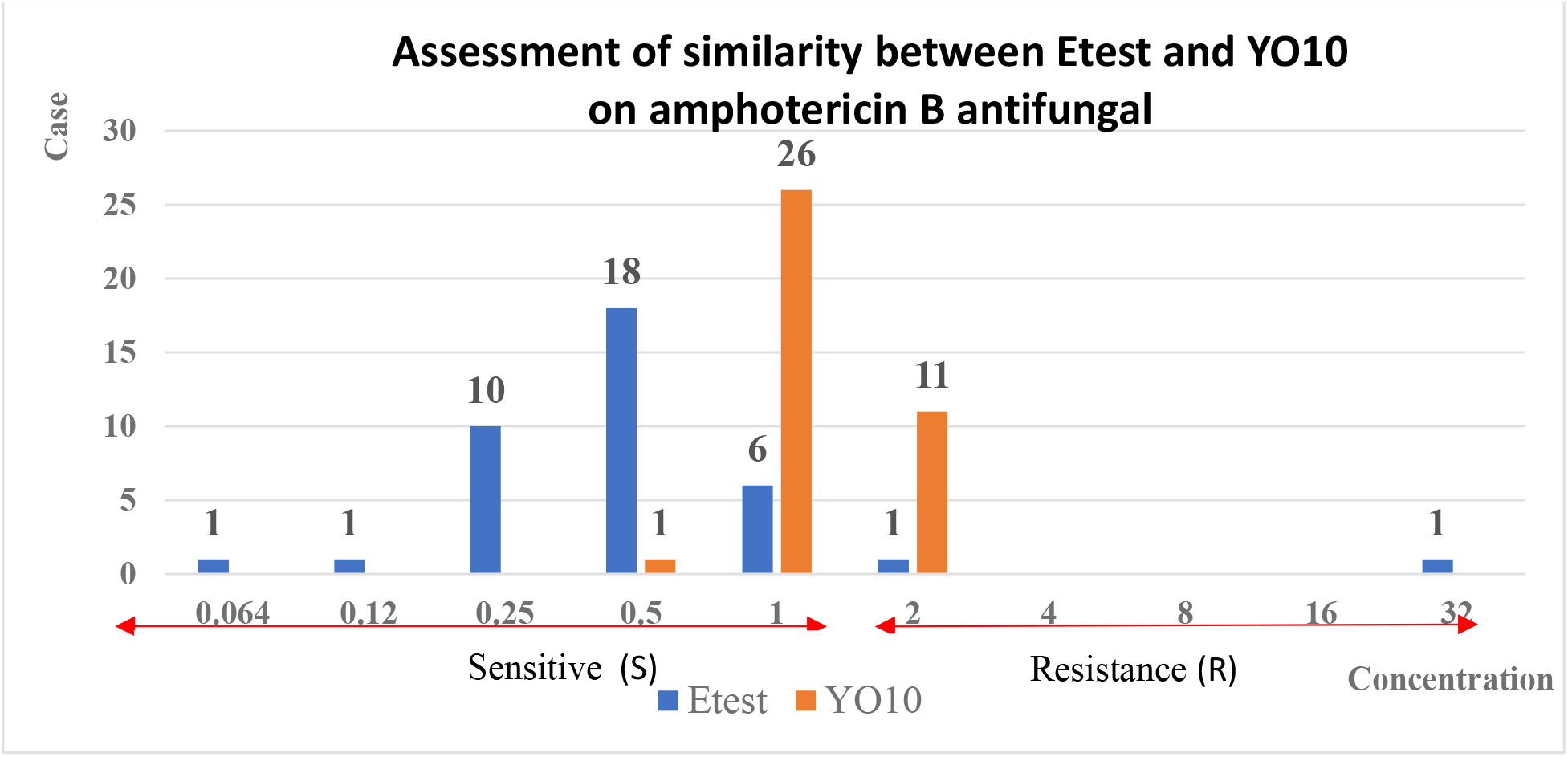
Assessment of similarity between Etest and YO10 on amphotericin B antifungal

In this study, it was noted that: MIC distribution was mainly from 0.25 - 1 μg/ml in Etest, and YO10 was 1 - 2 μg/ml; the similarity of MIC between the 2 methods was 2 samples accounting for 5.3%. MIC within ± 2 dilution limits between the 2 methods was 34/38 samples, accounting for 89.4%. MIC discordance between the 2 methods was 2/38 samples, accounting for 5.3%. Similar MIC distribution at 1 μg/ml (with 6 samples). The results of the antifungal test were similar in phenotype between the two methods: 29/38 samples, accounting for 76.3% (27 sensitive samples, 2 resistant samples), the results of the antifungal test were not consistent in phenotype between the two methods in 9/38 samples, accounting for 23.7% (9 samples had a sensitive phenotype in Etest but were resistant in YO10). When tested with the YO10 plate, there are 11 samples that fall right at the breakpoint (MIC = 2 μg/ml) and to be able to explain this difference, in the Clinical and Laboratory Standards Institute - M52 (CLSI - M52) guidelines for mycobacteria, the allowable precision sensitivity is ± 2 dilutions. [9] so 9 out of 11 samples with discordance at MIC = 2 μg/ml can be considered acceptable, so the discordance rate in terms of susceptibility phenotype (S) between the 2 methods is reduced to 0%. This is consistent with the study by Maria Siopi in 2023 [17], where the YO10 plate was 1 to 2 dilution orders higher (29%) and exceeded the “sensitivity”, thus resulting in significant discordance in terms of phenotypic susceptibility classification compared to the Etest MIC, with the Etest MIC of 1 μg/ml being only one dilution order lower than the US CDC breakpoint of 2 μg/ml, thus resulting in a discordance error between the two methods of 11%, while in this study it was 23.7%.

## Conclude

**Table 5.1:**
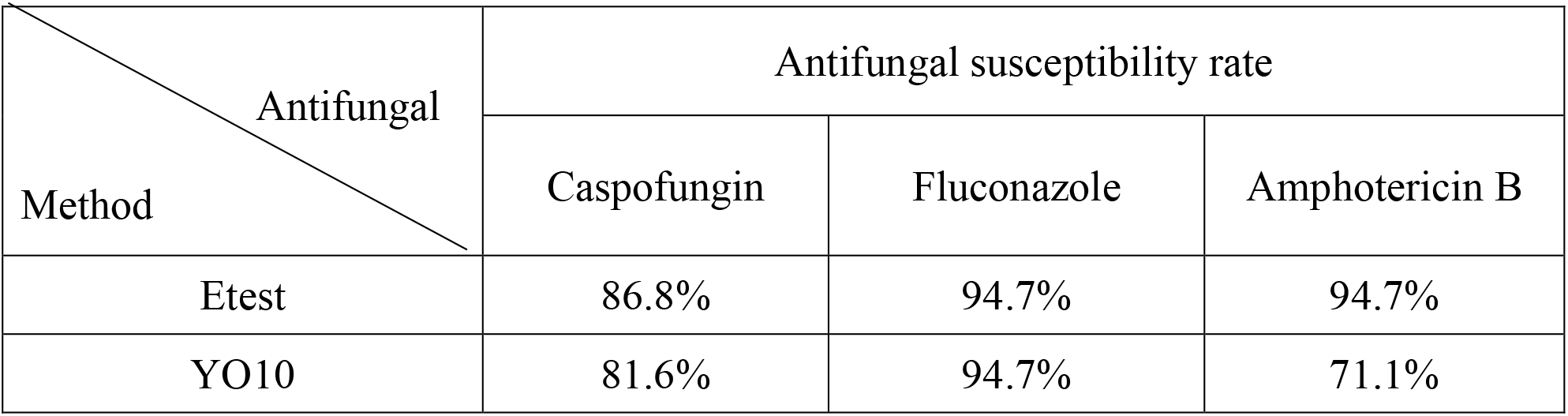
Antifungal susceptibility ratio between Etest and YO10 tray.

### Antifungal resistance rate of *C. auris* according to YO10 tray results

In this study, no cases of resistance to all three antifungal groups (azoles, echinocandins, polyenes) were recorded. However, there were still cases of resistance to 2 groups of antifungal agents accounting for 7.9% (3 out of 38 samples). Of these, 2 samples were resistant to both fluconazole and amphotericin B, and 1 sample was resistant to both caspofungin and amphotericin B. The rate of resistance to 1 out of 3 antifungal groups was 36.8% (with 14 samples out of 38 samples and randomly distributed among 3 antifungal groups).

**Table 5.2:** Evaluate the similarity between Etest and YO10 tray.

| Ratio \ Antifungal | Similarities between Etest and YO10 tray |  |  |
| --- | --- | --- | --- |
|  | Caspofungin | Fluconazole | Amphotericin B |
| MIC | 94.7% | 86.8% | 94.7% |
| Sensitive - resistant phenotype | 94.7% | 100% | 76.3% |

## Recommendation

### 6.1 Choose the appropriate method

In this study, we found that the similarity between the two methods is very high. Therefore, each laboratory needs to select an appropriate method to determine susceptibility for *C. aurris*. Priority should be given to choosing the method that is suitable for the existing conditions of the laboratory as well as economically for the patient and gives accurate results. Choosing the appropriate method and technique will contribute to providing clinical antifungal results accurately and quickly, increasing the cure rate, contributing to reducing hospital stay and reducing patient costs.

### 6.2 Antifungal resistance on future

Regarding the sensitivity of *Candida auris* to antifungal drugs, it is necessary to continue monitoring the sensitivity of *Candida auris* to antifungal drugs and take preventive measures in the future to avoid hospital outbreaks. Further analysis of the genome of *Candida auris* circulating in Vietnam as well as at Cho Ray Hospital is needed in future studies to determine the source of infection and compare it with other studies in the world. From the phenotypic results of susceptibility - resistance in antifungal assay, combined with molecular biological methods of gene sequencing of Candida auris to identify in - depth studies, creating conditions for future analysis and data sources in Vietnam as well as in the world.

